# Earbox, an open tool for high-throughput measurement of the spatial organization of maize ears and inference of novel traits

**DOI:** 10.1101/2021.12.20.473433

**Authors:** V. Oury, T. Leroux, O. Turc, R. Chapuis, C. Palaffre, F. Tardieu, S. Alvarez Prado, C. Welcker, S. Lacube

## Abstract

**Background:** Characterizing plant genetic resources and their response to the environment through accurate measurement of relevant traits is crucial to genetics and breeding. The spatial organization of the maize ear provides insights into the response of grain yield to environmental conditions. Current automated methods for phenotyping the maize ear do not capture these spatial features.

**Results:** We developed EARBOX, a low-cost, open-source system for automated phenotyping of maize ears. EARBOX integrates open-source technologies for both software and hardware that facilitate its deployment and improvement for specific research questions. The imaging platform consists of a customized box in which ears are repeatedly imaged as they rotate via motorized rollers. With deep learning based on convolutional neural networks, the image analysis algorithm uses a two-step procedure: ear-specific grain masks are first created and subsequently used to extract a range of trait data per ear, including ear shape and dimensions, the number of grains and their spatial organisation, and the distribution of grain dimensions along the ear. The reliability of each trait was validated against ground-truth data from manual measurements. Moreover, EARBOX derives novel traits, inaccessible through conventional methods, especially the distribution of grain dimensions along grain cohorts, relevant for ear morphogenesis, and the distribution of abortion frequency along the ear, relevant for plant response to stress, especially soil water deficit.

**Conclusions:** The proposed system provides robust and accurate measurements of maize ear traits including spatial features. Future developments include grain type and colour categorization. This method opens avenues for high-throughput genetic or functional studies in the context of plant adaptation to a changing environment.

## Introduction

Characterizing genetic resources and their response to the environment through accurate measurement of relevant traits is crucial to dissect the genetic bases of crop yield (Liang et al., 2016), and to tailor genotypes adapted to specific climatic scenarios (Tardieu et al., 2018). In maize, yield results from the number of grains and individual grain size, each of which has higher heritability than overall yield (Messmer et al., 2009; Peng et al., 2011), present different genetic architectures (Alvarez Prado et al., 2014; Amelong et al., 2015) and result from environmental conditions during different phases of the crop cycle, namely the vegetative and flowering period for grain number and the post-flowering period for individual grain weight (Gambín & Borrás, 2010). Sensitivities of grain number to soil water deficit, temperature and light are key parameters in the prevision of grain yield in a wide range of environments (Millet et al., 2019), thus requiring accurate phenotyping.

Examining the structure of the maize ear provides additional insights into deciphering the response of grain yield to environmental conditions. Indeed, the ear is composed of concentric rings of grains (cohorts) initiated simultaneously within each cohort but sequentially between cohorts (Messina et al., 2019) (Fig. 1 A-B). While the number of grains per cohort is a genetic trait largely independent of environmental conditions, the number of cohorts results from the response to climatic scenarios, with genotype-specific responses. Prior to flowering, suboptimal conditions reduce the number of grains via a reduction in the number of cohorts due to a reduced number of initiated ovaries (Moser et al., 2006). Abiotic stresses occurring at flowering result in localized ovary and grain abortion involving cohorts with delayed development (Oury et al., 2016) located preferentially at the ear apex, with aborted zone increasing with stress intensity (Fig. 1 C-D-E). Stress affecting pollination (pollen availability or viability) results in a wide variety of phenotypes characterised by incomplete cohorts and erratic cohort numbers (Fig. 1 F). Stress occurring beyond two weeks after flowering reduces grain size (Saini & Westgate, 1999). Therefore, a fine characterization of the spatial distribution along and around the ear of grain set/abortion and of grain and ear dimensions appears to be a relevant tool to reveal the response of genotypes to environmental scenarios.

**Fig. 1.**
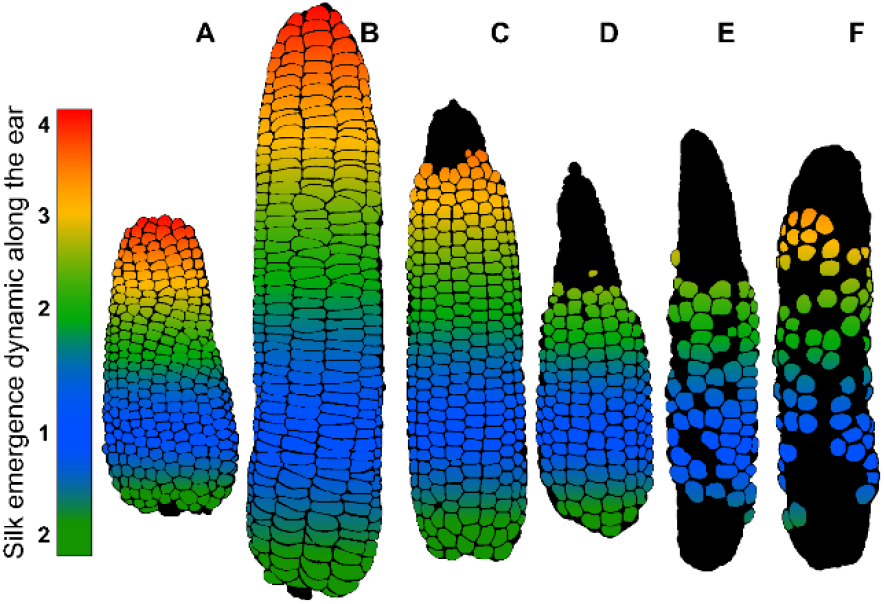
Spatial organization of grains reflecting the morphogenesis of the ear. The grains are arranged in rings and rows. Each ring corresponds to a cohort of organs with synchronous development, while a developmental gradient exists between cohorts depending on their vertical position along the rows. Floret cohorts are initiated sequentially at the ear apex. The oldest cohorts are located at basal positions and the youngest at apical positions. (A, B) Under optimal conditions, pollination and fertilization follow the order of silk emergence which is illustrated by colors: zone 1 cohorts (blue) are fertilized first, followed by zone 2 (green), zone 3 (yellow) and zone 4 (red). (C, D, E) Abiotic stresses occurring at flowering induce abortion that preferentially affects the youngest apical cohorts in zone 4, followed by the basal cohorts. (E F) Severe constraints affecting pollination (pollen availability or viability) result in a wide variety of phenotypes characterized by incomplete cohorts and erratic cohort numbers.

Ear phenotyping is still largely manual, time-consuming, costly, and subjective (Liang et al., 2016). Several methods have been developed to extract ear and grain characteristics from images (Chipindu et al., 2020; Makanza et al., 2018; Severini et al., 2011). They are usually based on manual or non-standardized acquisition involving either isolated grains after shelling (Liang et al., 2016; Makanza et al., 2018; Miller et al., 2017; Ni et al., 2019; Severini et al., 2011) or one side of the ear (Chipindu et al., 2020; Khaki et al., 2020, 2021; Kienbaum et al., 2021; Miller et al., 2017; Wu et al., 2020). Thus, the spatial distributions of grain presence/absence (grain set vs grain abortion) and grain traits along and around the ear is usually not, or only partly, considered.

Several techniques have been used for ear imaging, each providing different advantages and drawbacks. (i) Vertical positioning of the ear on a rotating axis allows imaging different sides of the ear at specific rotation angles (Grift et al., 2017; Warman et al., 2020). This method is efficient but only considers one ear at a time and requires time-consuming handling for ear positioning before imaging (1-2 min per ear). (ii) Portable imaging systems have been developed, directly threaded around the ear in intact field plants, imaging simultaneously all ear sides, allowing 3D reconstructions of the ear (Arvalis, 2018). This technique is affordable and avoids the need to harvest the ears, but involves limited throughput because of long ear handling time (husks removing, one ear at a time), while being subjected to various difficulties related to field conditions. Moreover, most of these techniques have been validated with ears from standard commercial hybrids with classical properties (ear and grain shape and colour, regular spatial organization), and therefore fail to provide reliable results for ears with non-regular patterns, a frequent characteristic under non-optimal environmental conditions (heterogeneity of abortion/set zones and grain dimensions, pest, and disease damage).

The aim of this study was to develop a low-cost and open-source system capable of producing automated, standardized, robust and reliable measurements of phenotypic traits of the ear, including the spatial distribution of grain traits along and around the ear. We tested it for genotypes with contrasting ear and grain shape and grain texture (e.g., Dent, flint, pop, waxy, flour). The method of image acquisition consisted of a custom box in which ears are placed horizontally. Motorized rollers are used to rotate the ears. In addition to being easy to setup and implement, this method is easily scalable for multiple ears at once by multiplying the number of rollers and cameras (Fig. 2).

**Fig. 2.**
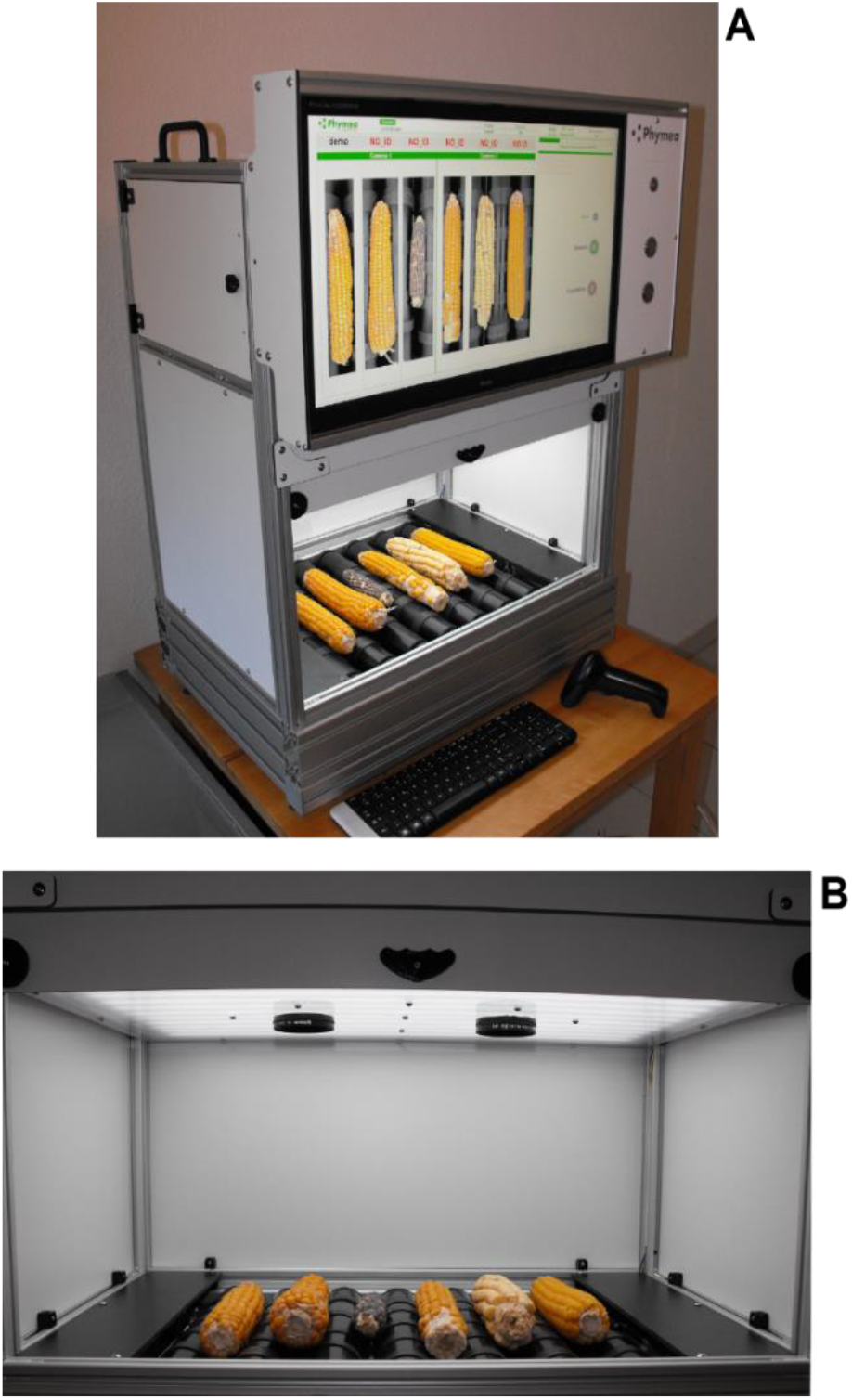
Pictures of the Earbox system. (A) The Earbox acquisition system, which allows the simultaneous acquisition of six maize ears on six sides via the rotation of the ear by motorized rollers. The identification of individual ears or ear lots is done by a keyboard or a barcode scanner. The system consists of aluminum profiles, compact laminates and assembly parts supplied by Elcom SAS or manufactured by Phymea Systems (CNC machining or 3D printing). The acquisition system is composed of two Raspberry Pi, each driving a Pi NoIR (V2.0) camera module, and a custom-made Arduino like board (ATMEGA 328P), to control the lighting and the two stepper motors (door and rollers) via two A4988 drivers. The master Rapsberry Pi (model 3 B+) hosts the main Python program, which centralizes all the functions of the system: the graphical user interface via the PyGame library, the control of the slave Rapsberry Pi (model B+) via SSH protocol, the communication with the Arduino like board for motor control, and the saving of the pictures to an external hard drive. (B) The Earbox system imaging cabin. Polarized lenses are added to the Pi Noir cameras. The lighting system consists of flexible LED strips in the visible (CRI 90) and infrared (940nm) wavelengths behind a frosted polycarbonate diffuser. Rubber strips are added to the rollers for optimal adhesion between the rollers and the ears.

We believe that this method will allow measurement of relevant traits in the context of plant adaptation to a changing environment and the enrichment of crop gene bank knowledge base (Law et al., 2011).

## METHODS

### A wide phenotypic diversity to test the robustness of the method

The set of ears used in this study was composed of 796 ears selected from two panels, a ‘biological diversity’ panel chosen to represent the diversity of phenotypes encountered in production contexts (Fig. 3 A), and an ‘environmental diversity’ panel, chosen to represent the phenotypes encountered in response to abiotic constraints (soil water deficit) (Fig. 3 B).

**Fig. 3.**
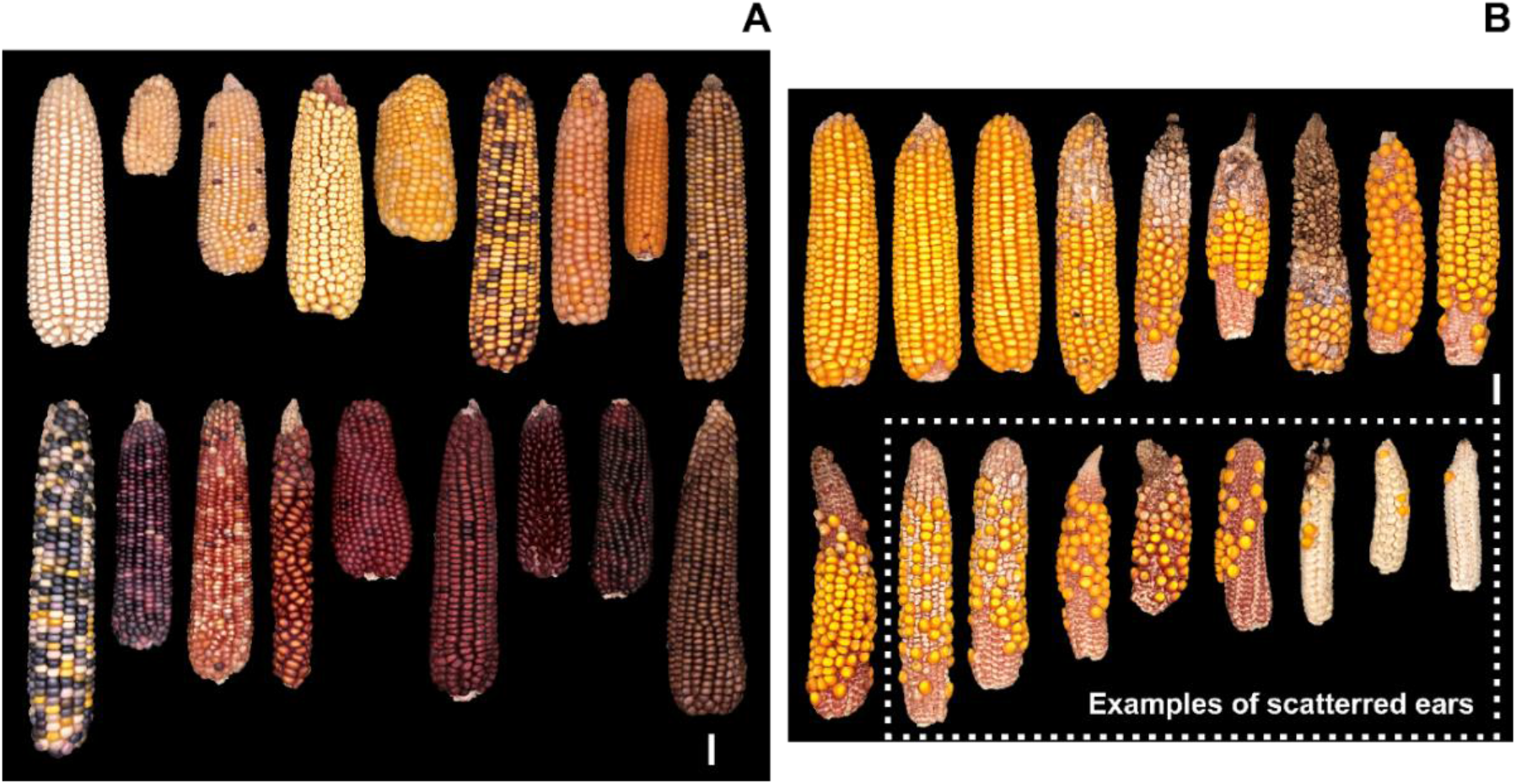
Representative ears from the biological and environmental diversity panels. (A) The biological diversity panel is mainly composed of lines and partially of non-commercial hybrids, with each ear often being a unique case. (B) The environmental diversity panel is composed of commercial hybrids obtained in an experimental context with biological treatments (well-watered and water deficits) and replications. These ears were selected to provide sufficient sampling to scan the full range of phenotypes encountered in a conservation context (A) and a production context (B) from optimal to near-zero (scattered ears). Dotted box, examples of scattered ears, which have incomplete cohorts all along the ear. White bar, 2cm.

The **Biological Diversity panel** represented 16% of the whole set, i.e., 126 ears. Selected ears were of various shapes (length between 3 and 24 cm and diameter between 2 and 5.5 cm). Grain colours were of all existing hues: white, yellow, orange, red, wine, pink, purple, blue, black, and brown, including heterogeneous ears with multiple colours and pearly, opaque, or translucent grains. Grain sizes ranged from 2 mm to 1 cm with variable shapes depending on their position along the ear, from perfectly round to dented or flint grains. Finally, the panel explored a diversity of grain spatial organization, with a range of number of cohorts and number of grains per cohort, and either regular or irregular grain organization along the ear.

The **Environmental Diversity panel** represented 84% of the set, i.e., 670 ears. First, a set of 431 ears was sampled from a field experiment (INRAE UE-DIASCOPE, France) under two water treatments: 321 under well-watered conditions (WW) and 110 under water deficit (WD). The remaining 234 ears were sampled in another experiment under WD treatment. For both experiments, water deficit conditions were triggered by stopping irrigation around 10-leaf stage while continuous irrigation was applied for the WW treatment. The combined variability in plant phenology and water treatments resulted in a wide range of ear phenotypes with various sizes and spatial distribution of fertile and aborted zones.

### A simple and low-cost image acquisition system

The ears were imaged with an automaton developed and assembled by Phymea Systems (www.phymea-systems.com - Montpellier, France). Individual ears are manually positioned in the system (Fig. 2), which acquires images stored in a generic hard drive. Images are uploaded to an independent analysis station where the associated software is installed for output retrieval. The automaton works in independent acquisition sessions to easily separate experiments, genotypes or varieties, and treatments. The ears or ear lots are individually identified by the keyboard or by a barcode scanner. The analysis software was developed to be flexible (retrieval of one or more phenotypic traits depending on user’s needs).

To minimize complexity and cost, the acquisition system was developed to be as simple and robust as possible. It consists of aluminium profiles, compact laminates and assembly parts supplied by Elcom SAS (Bourgoin-Jallieu, France) or manufactured by Phymea Systems (CNC machining or 3D printed) (Fig. 2A). The focus was on developing a flexible system, to be complexified in a second step: adapted for specific use cases, for example, to harvesting machines.

The system was designed to take multiple images of the ear via simultaneous rotations of all ears with motorized rollers. The acquisition system used in this study was set with 7 rollers for rotation and imaging of 6 ears (Fig. 2B). Rubber bands were added to the top of the rollers to properly drive the ears without slipping. Because the rollers have fixed dimensions (5.2 cm diameter) and positioning (1cm spacing), the theoretical ear rotation angle was calculated from the ear diameter and roller rotation angle and measured in practice by measuring the rotation of ears placed manually on the rollers. The measured and calculated angles fit strongly for the 4 ears tested, representative of the diversity of diameters in the whole ear panel (R^2^ = 0.98; Supp Fig. 1). We defined the number of images to be taken for each ear, thus the number of roller rotations, and a fixed roller rotation angle that ensured imaging of the whole ear circumference while minimizing acquisition time. The combination of 6 successive ear images with a roller rotation angle of 58° fulfilled these conditions for the range of 2 to 6 cm ear diameter (Supp Fig. 1) which exceeds the range encountered in both panels.

Developing a normalized method for analysing images regardless of ear or grain colour or shape required the use of near-infrared imaging. For this purpose, the system used Pi NoIR Camera v2 (Raspberry.org) driven by a Raspberry Pi to produce two types of images at two wavelengths: visible (RGB) and near infrared (IR, 940nm). Two sets of cameras and Raspberry Pi were necessary to ensure high resolution images of 6 ears at 6 angles and two wavelengths, for a total of twelve images, in less than 30 seconds. A custom Arduino-like board was developed to control both the lightning and the two stepper motors (doors and rollers). A master raspberry Pi was set to centralize all custom functions: host the main python program of the user interface developed with the PyGame python library, control the slave raspberry Pi via SSH protocol, control the Arduino board, and save the images to an external hard drive. The entire system was designed to be affordable and open source, and costs a total of 2500 € in equipment and hardware, excluding labour and development costs.

Ear peduncles at ear base were cut off prior to image acquisition, and husk-free ears were scrubbed and cleaned of silks and/or fungi with a brush so that all grains were accessible to the camera. A total set of 9492 images were taken from both panels.

### A combination of empirical segmentation and Deep Learning to build a robust routine workflow for Ear and Grain segmentation

RGB and IR images acquired from both panels of ears were first pre-processed (Fig. 4 A – Ground for deep learning) using conventional image analysis tools (dilate, open, close, gaussian blur and watershed) and merged to normalize the data for all ear and grain colours (Fig 4., Step 1), resulting in a pre-processed image set (4746 images: 6 images per ear for 791 ears, hereafter referred to as the ‘dataset’). The dataset images were then empirically segmented (Fig. 4, Step 2) and used to train a Deep Learning Neural Network (Fig 4, step 3). Finally, ear and grain phenotypic variables were retrieved for both ear panels (Fig. 4 B – Routine workflow). The ear masks were retrieved from the RGB images to estimate the ear phenotypic data (Fig. 4, Step 4). All images were then processed with the fitted neural network (Fig.4, step 5) and used to estimate grain phenotypic traits and characterize grain organization on the ear (Fig 4, Step 6).

**Fig. 4.**
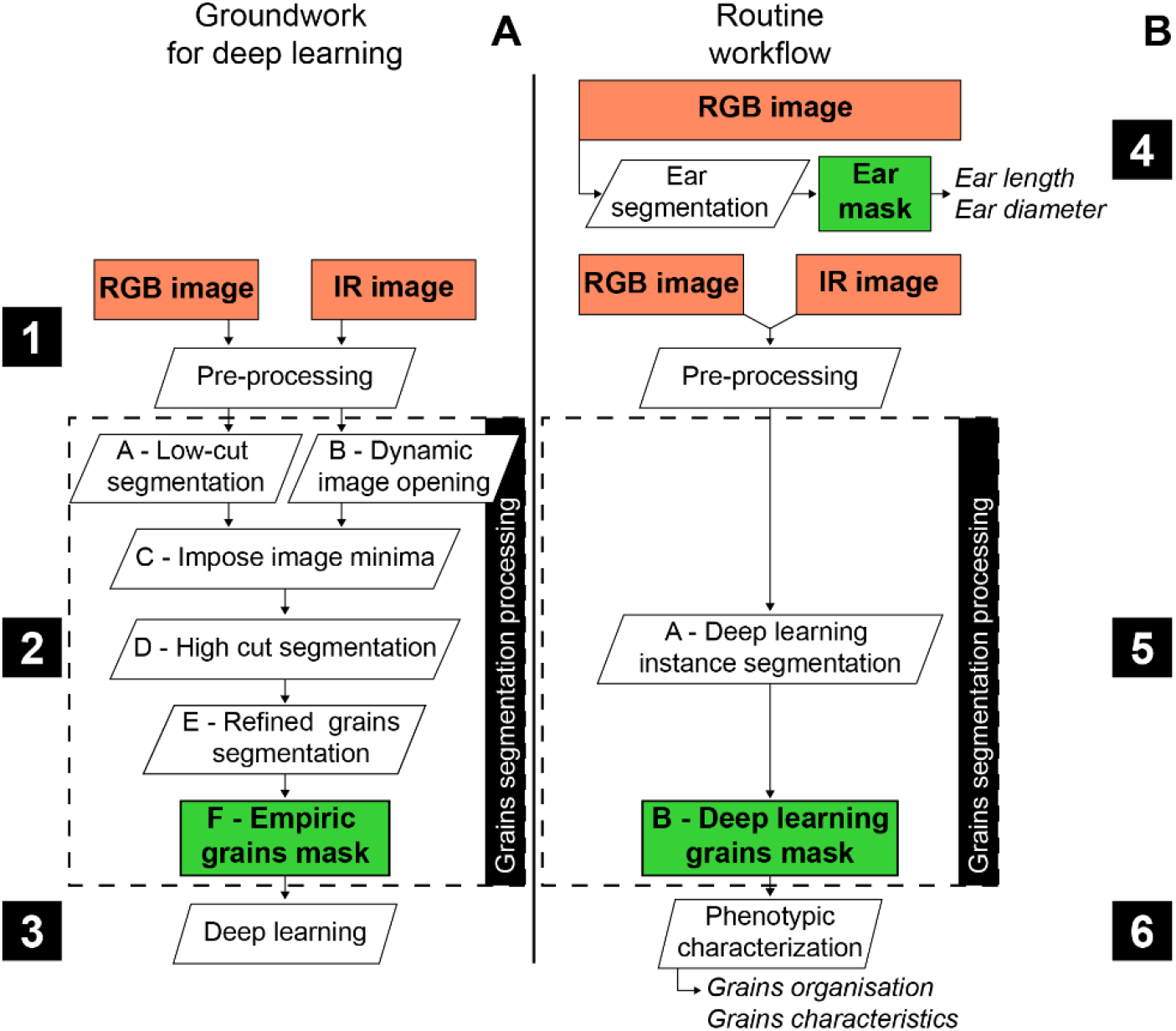
Workflows for training the neural network and generating phenotypic data. (A) Image processing workflow used to train the learning procedure (Groundwork for deep learning). (B), Image processing workflow used to produce phenotypic data (Routine workflow). Orange box, image acquisition. White box, image, or data processing. Green box processed images or data. Numbers in black boxes, steps of development, from first image acquisition to phenotypic data.

**A ground-truth dataset was built prior to any** analysis by manually segmenting grains from a set of images from 79 ears (10% of the dataset; not used to train the Neural Network). These ears were randomly selected from each category of grain colour and shape based on its frequency in the entire dataset.

**A preliminary step of empirical segmentation** was performed and used as an automatic annotation to drive deep learning iterations. The architecture of the empirical segmentation was developed to detect grains for a large portion of the dataset, so that all features can be learned and improved in Deep Learning run. RGB and IR images were processed with an algorithm developed by Phymea-Systems (Fig. 4, step 2) using only trial and error (using morphological image processing toolkits) to produce a grain mask that was precise enough to characterise the grains from images with various colours of grains and cob. The images were pre-processed with conventional image analysis tools to enhance their quality and then merged to retrieve complementary contrast and shapes. The cross-checking of the two pieces of information allowed a precise selection of the grains to produce an initial grain segmentation, which was in turn corrected by image analysis to refine the grain shapes and recover the over-segmented grains.

After this step, the processed images were sorted to assess the quality of the output. The output masks were scored by two independent individuals to evaluate the quality of segmentation with a score from 0 (bad segmentation) to 3 (good segmentation). Ears with uniform grain colour, strong colour contrast between cob and grain colour and non-scattered grains were mostly correctly segmented, with only minor problems. Most low scores were encountered for scattered grains (overly segmented), and for ears with similar grain and cob colour, which made it more difficult for the algorithm to distinguish.

**The Mask-RCNN neural network** was used for the Deep Learning training. It is commonly used as a framework for instance segmentation (He et al., 2018). It is a highly flexible, trainable framework that has been widely validated in many scientific domains, including plant science (Chipindu et al., 2020; Davis et al., 2020; Ganesh et al., 2019; Machefer et al., 2020; Wang et al., 2019; C. Zhang et al., 2020; J. Zhang et al., 2020). The entire deep learning (DL) framework was coded in Python 3 using TensorFlow for in-learning visualisation (Abadi et al., 2016). A set of data augmentation techniques were applied prior to learning. Each image was cropped into a set of 512 by 512 pixels elements on which a random number (between 0 and 2) of augmentation techniques were applied before being introduced into the model. The various augmentation transformations were retrieved from the ‘imgaug’ package (https://imgaug.readthedocs.io/en/latest/): flip up down, flip left right, 90 or 180- or 270-degree rotation, pixel value multiplication and gaussian blur. Deep Learning iterations involved 33 learning epochs with a cross-validation using 75 random images unused in the training dataset.

The neural network training (Fig. 4, step 3) included the following steps (Supp. Fig. 2). First, empirical masks with a mean score equal or greater than 2.5 (2076 images, 43.7% of the dataset, i.e., 346 ears) were used to train the neural network (Supp Fig. 2, step 3). The resulting DL1 masks outputs were corrected by ‘minor’ manual corrections (only ‘click on grains’ to add or remove mis detected grains) for 1926 images (92,8% of DL1 images, i.e. 321 ears) and ‘major’ corrections (adding grains and reshaping grains for ears with a large number of missing or mis-segmented grains and/or wrong shapes) for the remaining 150 images (7,2% of DL1, i.e. 25 ears – Supp Fig. 2, step 4). Second, a set of 72 images (12 ears, i.e., 1.5% of the dataset) from ears incorrectly segmented in the initial empirical segmentation, were manually corrected in the same way as the ‘major’ corrections seen above. Corrected images from DL1 and initial empirical segmentation were used in a second Deep Learning iteration (DL2 Supp Fig. 2, step 5) with 2148 images, i.e., 358 ears (45.3% of the dataset).

The ‘mean Average Precision’ (mAP) was used to estimate the quality of the Deep Learning output (Henderson & Ferrari, 2017) and calculated as defined by the latest evaluation’s techniques of the COCO dataset (Lin et al., 2015). The literature usually considers an algorithm to be highly efficient for mAP values of 0.4.

After several steps of learning, small input image correction, re-calibration of the neural network parameters, the resulting network with fitted weights (DL2) was used to extract segmented grains from all acquired images, i.e., the dataset (Fig.4., step 5).

### A routine workflow to access and validate phenotypic traits and their spatial distribution

Image analysis methods were applied on the segmented grains to extract phenotypic data for each trait of interest. To validate this methodology, the set of ears from both panels (791 ears) was also described, for each trait, by a unique observer to generate a set of manual measurements, to be compared to automatic measurements generated by the Earbox system (Supp Fig. 3). Most of the manual measurements were repeated 4 times around the ear circumference, averaged, and compared to the corresponding automatic measurements. The automatic measurements were repeated on each of the 6 images taken for each ear, and then averaged to produce phenotypic data at the ear scale.

**The ear dimensions and form were automatically acquired with the Earbox** from the segmented ear in each RGB image (Fig. 5A, B and C). All measurements were referenced to their spatial position according to the two axes of the image: the principal axis parallel to the ear length (vertical axis, starting from the bottom to the top of the ear) and the perpendicular axis (horizontal axis).

**Fig. 5.**
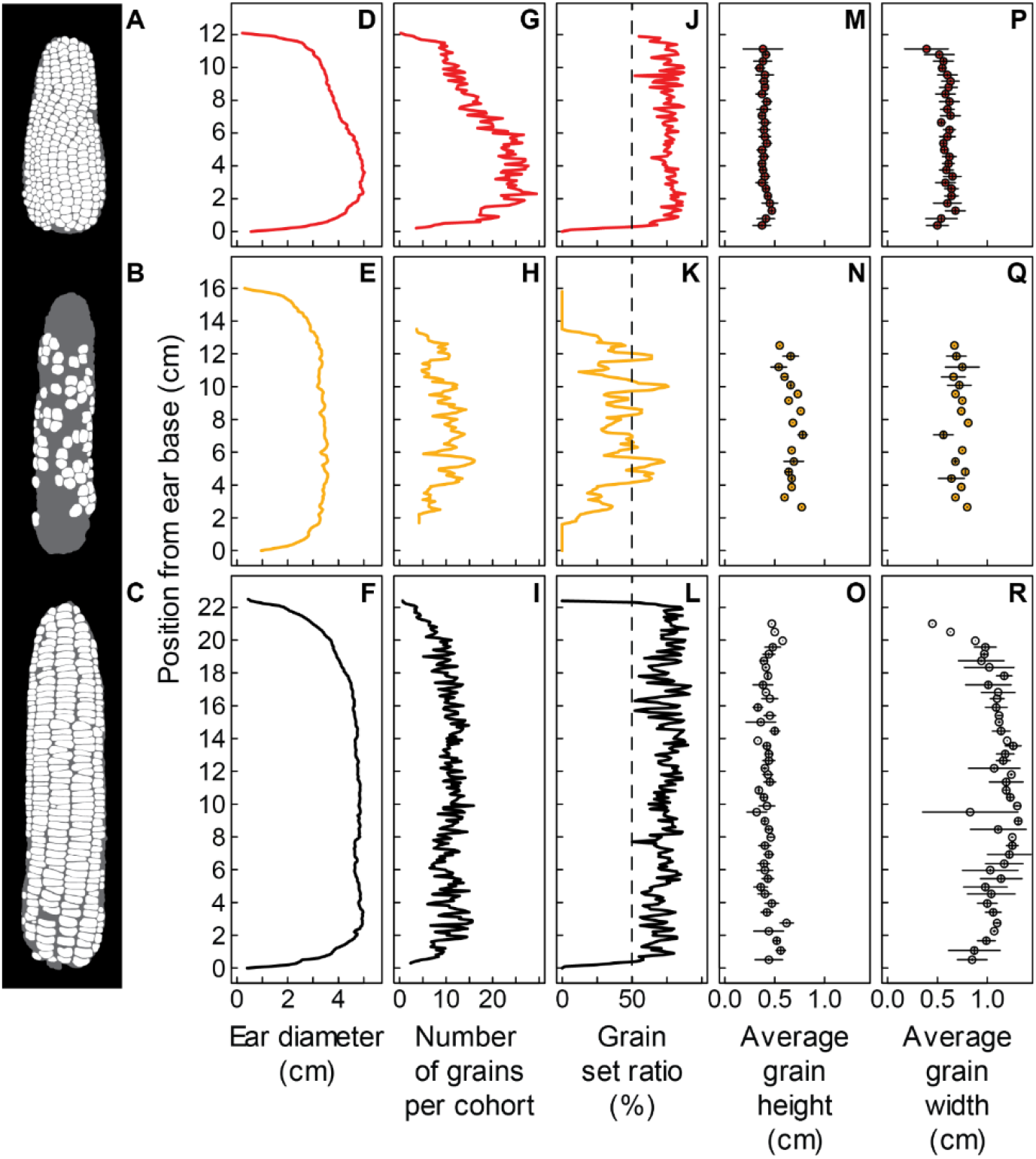
Raw phenotypic data measured with the Earbox analysis system. (A,B,C) Images from the grain segmentation process (output from the last deep learning iteration). (D,E,F) Measurements of ear diameter (x) for each pixel along ear length (y). (G,H,I) Results of the algorithm counting the number of grains per cohort (x) for each pixel along ear length (y). (J,K,L) Measurement of grain set ratio, the ratio of the number of pixels assigned to grains to the total number of pixels assigned to the ear (x) for every pixel along ear length (y), i.e. a measure of the percentage of the ear filled with grains. (M,N,O) Measurement of grain height (dimension along ear length) for each cohort of grain classified by the algorithm (one point = one cohort) according to its position along ear length (y). (P,Q,R) Measurements of the grain width (x) for each cohort classified by the algorithm (one point = one cohort) according to its position along ear length (y).

The ear mask was reduced to its centre pixel along the principal axis of the image (Supp Fig. 4A) to define the central axis of the ear. The *ear length* was calculated as the number of pixels of this central axis running from the bottom to the top of the ear. This method corrected for the twisting effect of irregular ear shapes. *Ear diameter* was measured at each pixel along the principal axis as the distance between the pixels of the ear contour (Fig. 5D, E and F).

Automatic measurements were tested by comparison with manual measurements. Ear length and maximum diameter were measured with a ruler and a sliding caliper (gauge), respectively. The measured length ranged from 3.4 cm to 23.8 cm and the maximum diameter from 1.9 cm to 5.5 cm, exploring similar variability for both panels.

**Grains were automatically counted with the Earbox** from the segmented grains in each image (Fig. 5A, B and C). Grain objects identified by segmentation were ‘shrunk’ either to a vertical line (one-pixel width) along the principal axis (Supp Fig. 4B) or to a horizontal line (one pixel height) along the perpendicular axis (Supp Fig. 4C). Horizontal distances between objects were corrected by considering each ear section *i* (one pixel height) along the ear axis as a circle of diameter *Diameter*_*i*_ (Supp Fig. 5). The mean distance *Dist*_*i*_ between two consecutive vertical lines was calculated at each position *i* along the ear axis.

The *number of grains per cohort* at position *i* was estimated by the Earbox as the ratio of the ear perimeter (π * *Diameter*_*i*_) to the distance *Dist*_*i*_ between contiguous grains (Fig. 5 G, H and I). The *number of cohorts* was estimated by the Earbox at each horizontal position perpendicular to the ear axis by counting the number of horizontal lines crossed from the bottom to the top of the ear (Supp Fig. 4C; Supp Fig. 6). The cohorts were incomplete on ear sides, and we considered the maximum observed value as the number of cohorts in the image (Supp Fig.6). The *number of grains per ear* was calculated by the Earbox from the number of cohorts and the number of grains per cohort measured in the 6 ear images and in the basal, median, and apical ear zones. It is derived from a composite calculation performed for each image by averaging: i) an over-estimator considering the number of grains per cohort in the median zone of the ear and the maximum number of cohorts, and ii) an under-estimator using information from both the number of grains per cohort and the mean number of cohorts in each third of the ear. The average of these two indicators was identified as the most relevant estimator.

**Manual measurements** related to grain organisation were performed manually to be tested against automatic measurements. The *number of grains per cohort* was counted at 3 positions along the ear by visually distinguishing a basal zone, a median zone, and an apical zone (Supp Fig. 3). It was compared to the mean number of grains per cohort averaged over the whole corresponding zone defined automatically *i*.*e*., basal third, median third, and apical third of the ear. The *number of cohorts* was counted on 4 sides of the ear and compared to the number of cohorts automatically calculated as an average over the 6 images (Supp Fig. 6). The *number of grains per ear* was counted with an automatic counting machine (*Contador: Seed counter – Pfeuffer GmbH (Quality control of grain and seeds)*, s. d.)) after removing them from ear cob. Because this measurement is destructive, it was only performed on a subset of the Environmental Diversity panel (257 ears), to keep enough ears intact to test and validate future updates and developments of the image analysis algorithm.

#### Grain dimensions were automatically measured with the Earbox

*Grain height and grain width* were calculated by fitting each segmented grain to a rectangle: grain height was defined as the fitted dimension along the axis of the ear and grain width as the fitted perpendicular dimension (Fig. 5 M to R). As mentioned above, the horizontal distance (grain width) was corrected to consider the circular shape of the ear sections (Supp Fig. 5).

Automatically measured grain dimensions were confronted to manual measurement. For this purpose, the height and width of 809 grains from the images of 9 reference ears from both panels were manually measured on the images generated by Earbox. We only considered grains located in the centre of the image to avoid distortion of grain widths, and performed a grain-by-grain comparison of dimensions by identifying each grain with its barycentre coordinates on the image.

**The spatial arrangement of grains in cohorts** was measured with the Earbox system by assigning each segmented grain to a cohort (Supp Fig. 7). To achieve this, the grains were scanned from the bottom to the top of the ear and classified according to their relative proximity, which depends on the mean size of objects (grains) in the image. More precisely, the algorithm starts with the lowest grain in the ear, checks the centroids of the grains at half the mean ear grain width and assigns the selected grains to the first cohort. Previously classified grains are removed for the remainder of the analysis and subsequent cohorts of grains are iteratively classified in the same way until no grains remain. This allowed grain dimensions to be plotted against cohort ranking (Fig. 5M to R) which is relevant to account for the developmental gradient along the ear due to morphogenesis (Fig. 1).

Automatic measurements of grains spatial arrangement were confronted to manual measurements. Each grain was manually assigned to a cohort by determining its rank along a row, from 1 at the ear bottom to *n* at the ear apex, at 3 positions around the ear.

#### Finally, an automatic method for abortion zones characterization was developed

Grain masks allowed discriminating grain pixels from pixels outside the grains but within the ear contours. The latter were considered as corresponding to aborted zones. The *grain set ratio* was calculated for each ear section *i* (one pixel height) along the ear axis as the ratio of the number of grain pixels to the number of pixels of ear diameter at that section. It was smoothed by a running average over 2% of the total vertical pixels to make it less sensitive to high variations (Fig. 5J, K and L). At the ear level, we considered the zones with grain set ratio greater than 50 % as fertile zones and the others as aborted zones. When no fertile zone was detected, its length was set to 0 and the apical and basal aborted zones were each set to half of the total ear length.

The automatic method of abortion characterization was tested against manual measurements. The dimensions of the aborted and fertile zones were visually positioned and measured manually with a ruler to represent the approximate positions at which the cohort abortion rate was greater (aborted zone) or lower (fertile zone) than 50%.

## Results

### A high-quality segmentation allowed for reliable trait measurements

The masks computed with the trained neural network provided a standardized and reliable method for ear and grain segmentation on the whole set of acquired images. The mean average precision metric (mAP) used to assess the quality of segmentation with manually segmented images was 0.4 for Empirical Segmentation (ES) masks and 0.52 and 0.55 for Deep Learning DL1 and DL2 results, respectively. The ES masks had an acceptable value above the standard threshold, while both DL1 and DL2 iterations improved the indicator. In addition to validating the quality of the segmentation, these results also show an overall improvement at each step, highlighting their importance in the method. The high-quality segmentation obtained for a wide diversity of ear and grain phenotypes allows the production of comparable, standardized, and automatic data for both studied panels, composed of ears as different as small horny strawberry-shaped, and dented commercial hybrids.

The use of Neural Networks for Deep Learning is a powerful tool for image analysis, commonly used for plant phenotyping, both for morphological measurements and feature classification. The downside of our method is its requirement for annotated images that are difficult to acquire and analyse: they involve time-consuming annotation work, often performed manually. To overcome this, we chose an empirical approach using simple image analysis tools to produce a set of automatically annotated ear images with little time-input (avoiding manual annotation of the whole set of pre-processed images) to be used for deep learning iterations. The resulting empirical masks were used to train a Neural Network. Indeed, this is a direct and efficient mean to synthesize the best information while potentially improving the outputs. Furthermore, it provides a straightforward and efficient way to improve the analysis system if needed, by adding information through new segmented masks from previously unaccounted-for diversity.

### Ear dimensions and shape, fertile and aborted zones

The automatic measurements of ear length and ear diameter were accurate. The linear regressions with the manual measurements closely fit to the bisector for both variables: respectively R^2^ = 0.99, RMSE = 0.44cm for ear length; and R^2^ = 0.97, RMSE = 0.15cm for ear diameter (Fig. 6A and 6B). The small differences may be due to differences in methodology: the automatic algorithm measured the ear length with the joined line of the central pixels of the ear (usually not a straight line) while the manual measurement was a straight-line measurement. Since most ears are curvilinear, the automatic measurement appears to be more accurate in describing the diversity of ear shapes. For the same reasons, the algorithm might also be more accurate in determining the maximum ear diameter, which was inferred via an algorithm, rather than visually. Moreover, the accurate measurements of the algorithm at each pixel along the image (Fig. 5D, E and F) provide access to new variables describing ear shape, such as ear diameter along the ear and centreline curvature, to be investigated and explored for new avenues of comparison between phenotypes and varieties.

**Fig. 6.**
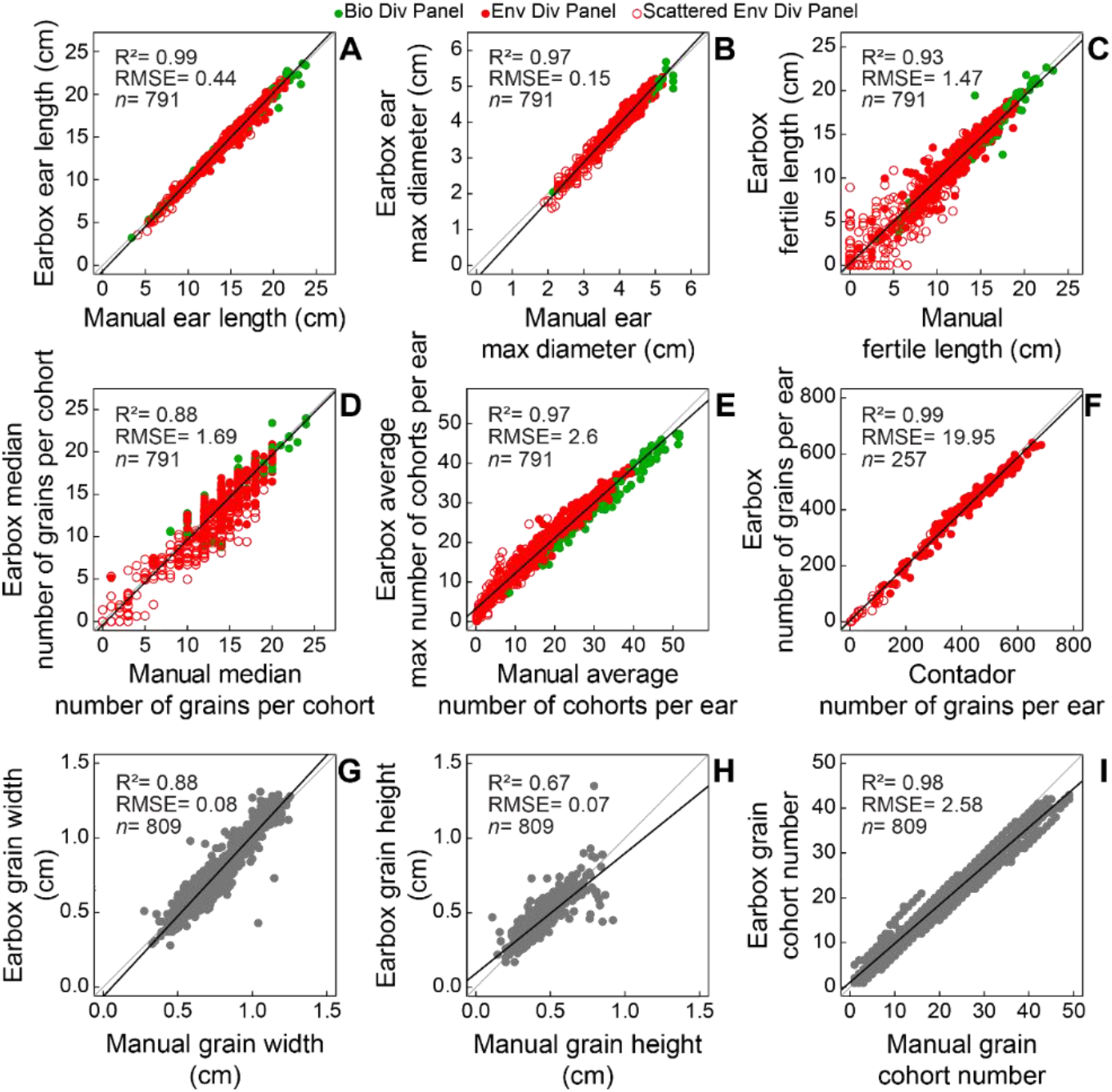
Comparison of Earbox (y) and reference (x) data. (A) Ear length in centimeters. (B) Maximum ear diameter in centimeters. (C) Length of the fertile zone in centimeters. (D) Number of grains per cohort in the median third of the ear. (E) Average number of cohorts per ear, the mean of 4 observations around the ear (x), the average of the maximum number of cohorts per ear of 6 images of the ear (y). Graphs A-E include all 791 ears from both panels. (F) Number of grains per ear measured automatically by the Earbox (y) and manually with an automatic grains counter (x) for 257 ears selected from the environmental diversity panel. (G) Grain width in centimeters, measured along the axis perpendicular to the ear. (H) Grain height in centimeters, measured along the main axis of the ear. (I) Assignment of a cohort number to each grain. Data presented in G-I correspond to a set of 809 grains measured and classified automatically by the Earbox (y) and measured manually on the acquisition images (x) for comparison. Green dots: ears from the biological diversity panel; red dots: ears from the environmental diversity panel. Empty red dots: scattered ears of the environmental diversity panel (Fig. 3 B). Grey dots: grain dimensions (one dot = one grain). Grey line: bisector line. Black line: linear regression of the data. R^2^: correlation coefficient between x and y values, RMSE: root mean square error, *n*: number of observations in each graph.

Promising results were obtained towards a standardized way of characterising maize ear abortion. Comparison of automatic and manual data for fertile and aborted zones (fertile zone: Fig. 6C; apical and basal abortion zones: Supp Fig. 8C and 8D) indicates good agreement between them, especially for the fertile zone length (R^2^=0.93, RMSE = 1.47cm; Fig. 6C). A discrepancy appears in the case of scattered ears (empty dots; Supp Fig. 8C and 8D) for which a precise visual positioning of the zones is difficult because the grains are randomly scattered along length and circumference of the ear (Supp Fig. 6). In these cases, the automatic method is probably more relevant, as it uses a common and accurate rule for all ear types. Moreover, characterization at each vertical pixel provides information on the spatial distribution of abortion and grain set along the ear (Fig. 5J, K and L), inaccessible by conventional methods.

### Grain counting

The spatial organisation of grains along the ear measured with the automatic method was both accurate and trustworthy, even for scattered ears. First, for the number of grains per cohort, the manual and automatic estimators were highly correlated in the median (R^2^=0.88, RMSE = 1.69 grains; Fig. 6D) and basal zone (R^2^=0.87, RMSE = 2.07 grains; Supp Fig. 8A), and less correlated in the apical zone (R^2^=0.68, RMSE = 3.43 grains; Supp Fig. 8B). Most of the discrepancies are due to scattered ears (empty dots in Fig. 6 and Supp Fig. 8) *i*.*e*., ears with incomplete cohorts, making the cohort identification uncertain and, as a result, counting their grain number difficult (Fig. 3B; Supp Fig. 3, scattered ear). Earbox data tend to be more objective and closer to the true average number, as they incorporate information from the entire apical zone, whereas manual estimates can be considered more subjective where abortion was high (empty dots in Fig. 6D and Supp Fig. 8A and B). In addition, the Earbox data provide access to the vertical distribution of this variable (Fig. 5G, H and I).

Manual and automatic measurements were also highly correlated for the number of cohorts (Fig. 6E, R^2^ = 0.97, RMSE = 2.6 cohorts), even for scattered ears.

The number of grains per ear was highly variable, ranging from almost 0 to about 700 grains per ear (Fig. 6F). The Earbox estimator of the number of grains per ear, which considers the number of cohorts and the number of grains per cohort, was highly correlated with the Contador measurement (Fig. 6F; R^2^ = 0.99, RMSE = 19.95 grains). These results validate the potential of the whole system to be used under both optimal and constraining conditions and for the study of the determinism of grain number in maize and its response to the environment.

### Grain dimensions and spatial positions: a potential framework for studying developmental gradients

The method adequately estimates grain dimensions (Fig. 6G and H), ranging from 0.3 to 1.3cm wide (Fig. 6G) and 0.1 to 0.9cm high (Fig. 6H) in both panels. The correlation between manual and automatic measurements was indeed high for grain width (R^2^=0.88, RMSE = 0.08cm; Fig 6G), and slightly lower for grain height (R^2^=0.67, RMSE = 0.07cm; Fig 6H). The slight differences may be due to the narrower range of variation observed for grain height versus grain width in the training dataset, which can be easily addressed with further Deep Learning iterations with suitable datasets. Most of the noise comes from isolated grains on scattered ears that tend to have a more circular shape when space is available around them (examples Fig. 3B), which could be easily resolved with an increase in the proportion of data or an individual training for scattered ears. Nevertheless, the results indicate that the method correctly positions the grain barycentre and properly captures shape variations between and within ears.

Automatic and manual measurements were consistent in assigning a cohort number to each grain, *i*.*e*., its vertical positioning along ear rows (Fig. 6I). Manual and automatic grain cohort numbers were highly correlated in the 809-grains sample set (R^2^=0.98, RMSE = 2.58 grains), indicating that the grains were properly located in the spatial organisation of the ear.

Thus, the system was able to characterize and discriminate a large variability in grain dimensions (width and height) and shapes (width/height ratio) among the studied ears (Supp Fig. 9), potentially allowing a reliable characterisation of genotypes based on these traits. In addition, by gathering grains into cohorts with synchronous development, the method gives access to the distribution of grain dimensions along developmental gradients. These distributions differ among ears: they are almost flat, decrease at different rates, or display a maximum at different vertical positions (Supp Fig. 9). Since developmental gradients are relevant to ear morphogenesis, they provide a framework for study and analysing the determinism of grain dimensions and, consequently, grain filling and weight.

### The spatial distribution of abortion reveals plant response to stress

The grain set ratio (GSR) emerged as a synthetic trait characterizing the response to environmental scenarios (Fig. 7). GSR profiles, i.e., grain set ratio as a function of vertical position along the ear axis, were established for the environmental diversity panel grown under contrasting soil water availability during flowering (665 ears). We performed an a priori-free analysis of all these profiles, using a clustering algorithm (k-means). Vertical positions were normalized by ear length to be independent of absolute ear length, and averaged for all sides of each ear. All curves were processed using Euclidean distance matrix and ward method to generate a cluster tree synthetizing the similarities between ears and finally discriminating ears into 5 clusters with increasing intensity of abortion from cluster 1 to cluster 5 (Fig. 7A, D). Cluster 1 coincided with ears with no or limited abortion. Abortion was limited to the apical zone of ears in cluster 2 and extended to the basal zone in cluster 3. The apical and basal aborted zones were wider in cluster 4, whereas they extended to the entire vertical profile of the ears from cluster 5. The progress of abortion from cluster 1 to cluster 5 followed the reverse order of silk emergence, which is reported in the literature as the main predictor of ovary/grain abortion frequency in response to constraints during flowering (Fig. 1 and Oury et al., 2016). The distribution of well-watered and water-stressed plants among clusters also indicated an increasing impact of stress from cluster 1 to cluster 5. Well-watered plants mainly belonged to cluster 1, and barely to clusters 2 and 3, whereas water-stressed plants mainly belong to clusters 3 to 5 (Fig. 7B). Moreover, the average ear length decreased from cluster 1 to cluster 5 whereas it was not considered as a factor to sort clusters (Fig. 7C).

**Fig. 7.**
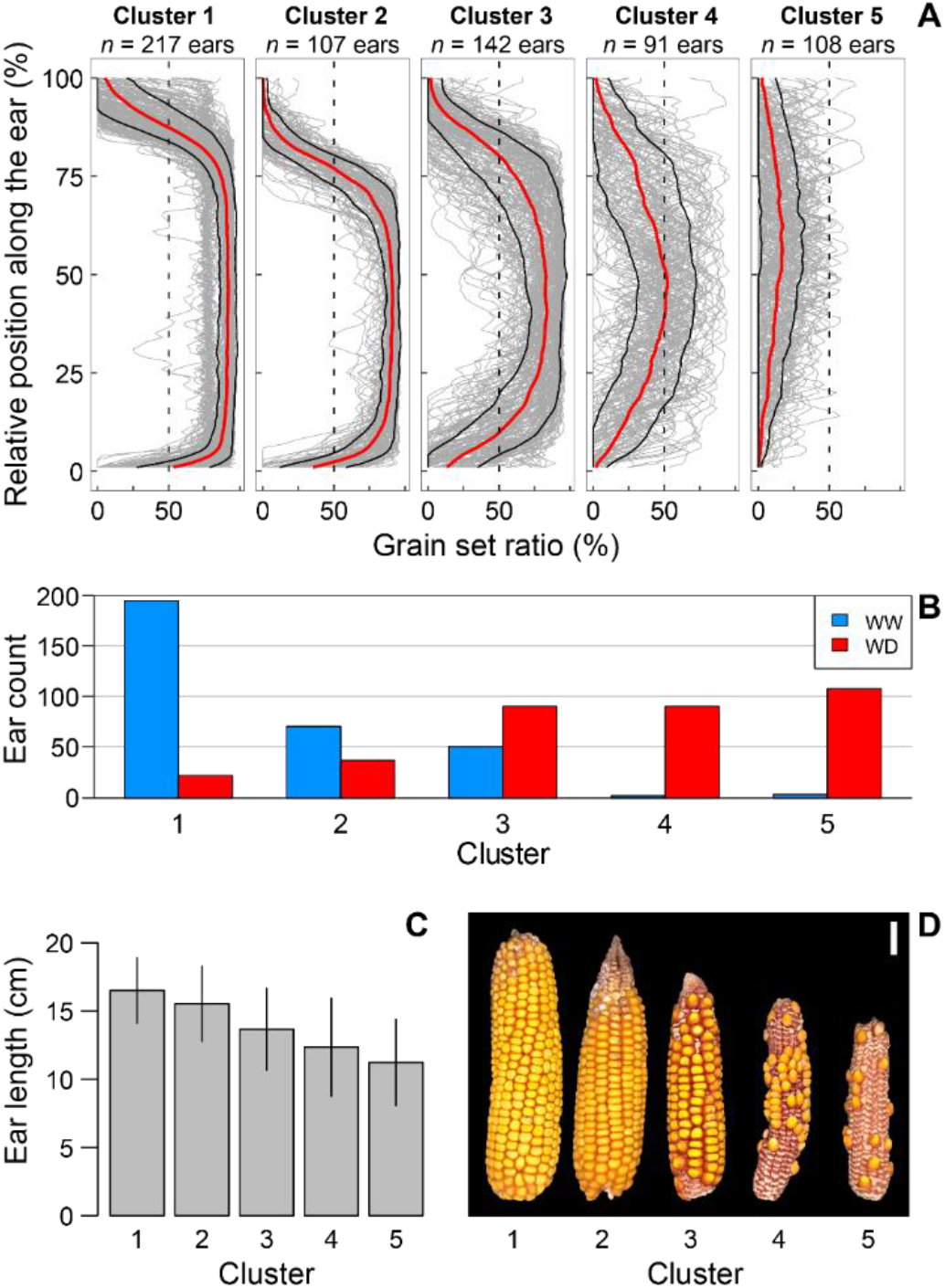
Clustering of grain set ratio reflecting silk emergence dynamics. (A) Grain set ratio (%) as a function of relative position along the ear (%). Ear positions are divided by ear length for easier comparison between ears. Grey lines: one line for each ear in the cluster. Red line: smoothed mean value of each cluster. Black lines: standard deviations from the mean line of each cluster. (B) Proportions of ears from each treatment (WW = well-watered, WD = water deficit) in the respective clusters (1 to 5). (C) Bar plot representing mean values of ear length in centimeters for each cluster, error bar = standard deviation. (D) Phenotypes of ears representing each cluster.

## Discussion

### A robust phenotyping pipeline to evaluate biological resources, complementary to existing procedures

The phenotyping pipeline developed and presented in this study (hardware and software) was able to accurately characterize, independently of the colour, shape, or transparency of grains and ears: the shape and dimensions of the ear, the number of grains and their spatial organisation, and the dimensions of the grains along the ear. The data were very similar to conventional manual data, with a much lower acquisition time. In addition, the system provides new traits, inaccessible by conventional methods, especially grain dimensions as a function of the grain cohort number, relevant to ear morphogenesis, and the distribution of abortion frequency along the ear, relevant to plant response to stress. Analysis of the genetic bases of these traits could enlighten the role and regulation of crucial genes determining ear phenotype and grain organisation, as well as their response to the environment. This could lead to new breeding traits responding to climatic challenges, which could be used in marker-assisted selection.

Adding to our system an automated and standardized calculation of standard qualitative descriptors of maize cultivars, such as the conicity of the ear or, for the grains, their shape, type, colour, or organization on the ear (GEVES or UPOV technical documentation) would only require simple general statistical classification methods based on machine learning or deep learning.

Our methodology is also complementary to other methods used for varietal description, such as cross-sections or ear deseeding, to characterize the cob, or the morphology of the grains and their physiological characteristics (Baye et al., 2006; Spielbauer et al., 2009), involving robotics for ear and grain handling (« Seed Analysis Automation », s. d.). Its relative simplicity and flexibility allow easy adaptation of the ear processing line to integrate the Earbox phenotyping solution before ear deseeding and grain phenotyping.

### Contribution to the analysis of adaptation/tolerance to environmental scenarios in combination with crop models

The new features obtained by the phenotyping pipeline open new avenues in the characterisation of maize grain yield formation in response to genetic and/or environmental factors. In particular, the spatial distribution of the grain set ratio appears to be a marker of the dynamics of silk emergence and ovary/grain abortion, a major component of the plant’s response to environmental scenarios(Oury et al., 2016; Tardieu et al., 2018). Consistent with the literature (DeBruin et al., 2018; Liu et al., 2020; Oury et al., 2016; Shen et al., 2018) our results (Fig. 7) suggest that the measurement of this variable potentially provides a high-throughput proxy for the complex processes involved in ear morphogenesis (rate and number of ovary initiations, growth rates of silks and pollen tube, development of the husks). This would greatly facilitate ecophysiological studies of the mechanisms determining yield components and their response to the environment.

Yield losses in maize are most pronounced when stress occurs around flowering (Campos et al., 2006; Claassen & Shaw, 1970) affecting grain number determination. Reproductive failure has different faces and can manifest as ear barrenness, incomplete ear pollination due to lack of pollen, and grain abortion (Cárcova et al., 2003; Cárcova & Otegui, 2001; Moss & Downey, 1971; Oury et al., 2016). The effects of the timing of stressful conditions and the pattern of zygote development along the ear row (successive cohorts) determine the nature of the ear phenotype associated with reproductive failure (Messina et al., 2019). In this sense, the ear phenotype can tell whether the crop experienced a stressful scenario and give some clues about the timing of the stressful condition. However, its implementation in plant genetics is difficult or impossible due to the size of studied panels of genotypes and/or to the number of traits resulting from stress x phenology combinations. The Earbox system presented in this study answers this bottleneck by providing standardized and high-throughput traits measurement. Furthermore, such a development paves the way towards new analyses to understand the interaction between genotype and environment in the context of water deficit for maize. While many efforts have been made to study the dynamics of water deficit related to phenology, (Chenu et al., 2013; Harrison et al., 2014), less progress has been made to study its genotypic variability under drought scenarios (Messina et al., 2019). As such, additional investigations provide insights to link the observed phenotype at harvest with events occurring at flowering, thus helping in the identification of varieties best adapted to a specific water deficit scenario.

## Conclusion

The system developed and presented in this study is a scalable system providing an accurate, robust, and reliable way to extract precise measurements from maize ear images, including spatial features of grain organization. This work illustrates, like many others (Bagley et al., 2020; Czedik Eysenberg et al., 2018; Gaggion et al., 2020; Pearce, 2020; Salter et al., 2020), the possibilities and the efficiency that open-source technologies and low-cost electronics now offer to plant science. They make accurate phenotyping accessible to everyone. In the case of the Earbox, even research structures with limited resources, farmer cooperatives, or multi-site research projects (limited by multiple observers and non-standardized methodologies), can claim reliable and reproducible ear phenotyping data with a system that can be easily modified to be integrated into complete ear and grain processing chains. For example, cameras can be replaced for higher resolutions or multispectral acquisition for characterization of grain physiology (Caporaso et al., 2018; Chu et al., 2020; Pang et al., 2020; Türker-Kaya & Huck, 2017; C. Zhang et al., 2020; J. Zhang et al., 2020; Zhou et al., s. d.). Additional steps of deep learning would probably be sufficient to develop a method for the recognizing and classifying of maize diseases(Hobbs et al., 2021; J. Zhang et al., 2020), or for characterising early grain development, by processing immature ears and grains a few days after flowering. Finally, the results of this work pave the way for future development of tools for inflorescence phenotyping of other crops, such as wheat and sunflower, for which the present system will be adapted.

## Authors contributions

**Conceptualization**^**\***^: Vincent Oury, Timothé Leroux, Olivier Turc, Sébastien Lacube, Carine Palaffre, Romain Chapuis, Claude Welcker

**Project supervision:** Vincent Oury

**Project administration:** Vincent Oury, Sébastien Lacube, Olivier Turc, Timothé Leroux

**Biological resources & experimentation:** Carine Palaffre, Romain Chapuis, Claude Welcker

**Data measurements & gathering:** Vincent Oury, Romain Chapuis, Carine Palaffre, Timothé Leroux, Sébastien Lacube

**Hardware-development:** Vincent Oury, Timothé Leroux

**Software-development:** Vincent Oury, Sébastien Lacube, Timothé Leroux

**Deep Learning implementation & training:** Timothé Leroux

**Formal analysis & investigation:** Sébastien Lacube, Vincent Oury, Timothé Leroux, Olivier Turc

**Data visualization:** Vincent Oury, Sébastien Lacube, Olivier Turc

**Writing – review & editing:** Sébastien Lacube, Vincent Oury, Olivier Turc, François Tardieu, Santiago Alvarez Prado, Timothé Leroux, Claude Welcker

^*\**^*Authors are listed in the order of contribution for each item*

## Acknowledgments

We thank all the staff of the Station Expérimentale de Mauguio & Saint-Martin-de-Hinx for their special contribution to grow, harvest and provide access to the biological diversity used in the dataset of this study. We particularly thank the French INRAE agents, especially Serge Malavieille, for their patience during the testing of our prototypes, their feedback and their valuable knowledge that allowed us to successfully complete this project. A special thanks to Christian Fournier (INRAe LEPSE) for his helpful advice to put in place the first steps of the empirical segmentation. Special thanks to Romain Chapuis and Olivier Turc for investing time in the project from beginning to end. Finally, we thank Joubin Ezami for his patience and his valuable contribution for the 3D modeling and the production of the imaging system.

## Notes

### Competing Interest Statement

The authors have declared no competing interest.

