## Supplemental figures for "Earbox, an open tool for high-throughput measurement of the spatial organization of maize ears and inference of novel traits"

### SUPPLEMENTAL MATERIALS

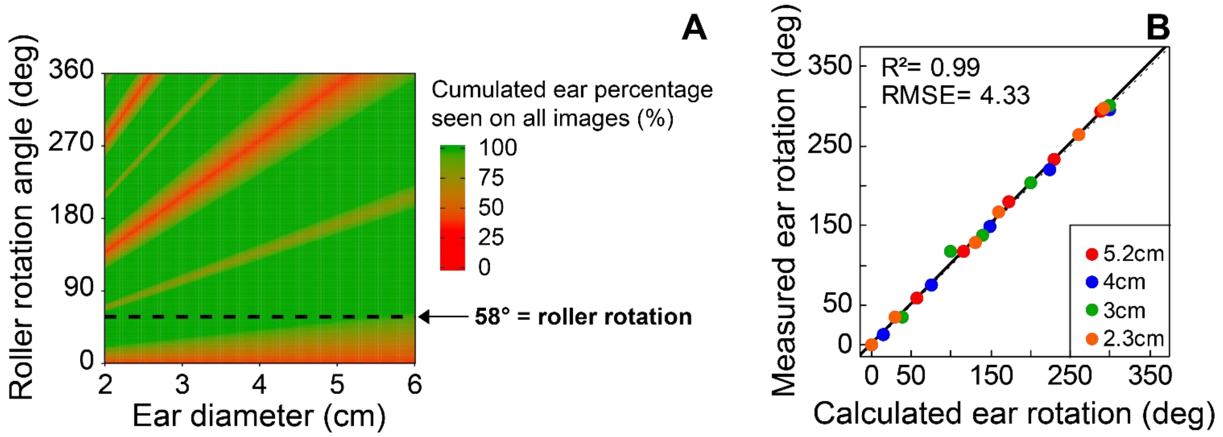


**Supplementary Fig. 1. Choice of ear rotation angle based on Earbox specifications.** (A) Heatmap of the theoretical cumulative ear percentage seen in all 6 images for roller rotation angles between 0° and 360° and diameters between 2cm and 6cm. Values are calculated assuming 120° of the ear circumferences is captured in each image. The dotted line represents the selected roller rotation angle, for which no critical diameter is encountered, capturing 100% of the information from each ear. (B) Match between theoritical and measured rotation angle for a set of ears chosen to represent diameter diversity in maize. Dots: measured and theoritical rotation angles. Colors: ear diameters. Dotted line: y = x. Solid line: linear regression. $R^2$: correlation coefficient between x and y values, RMSE: root mean square error.

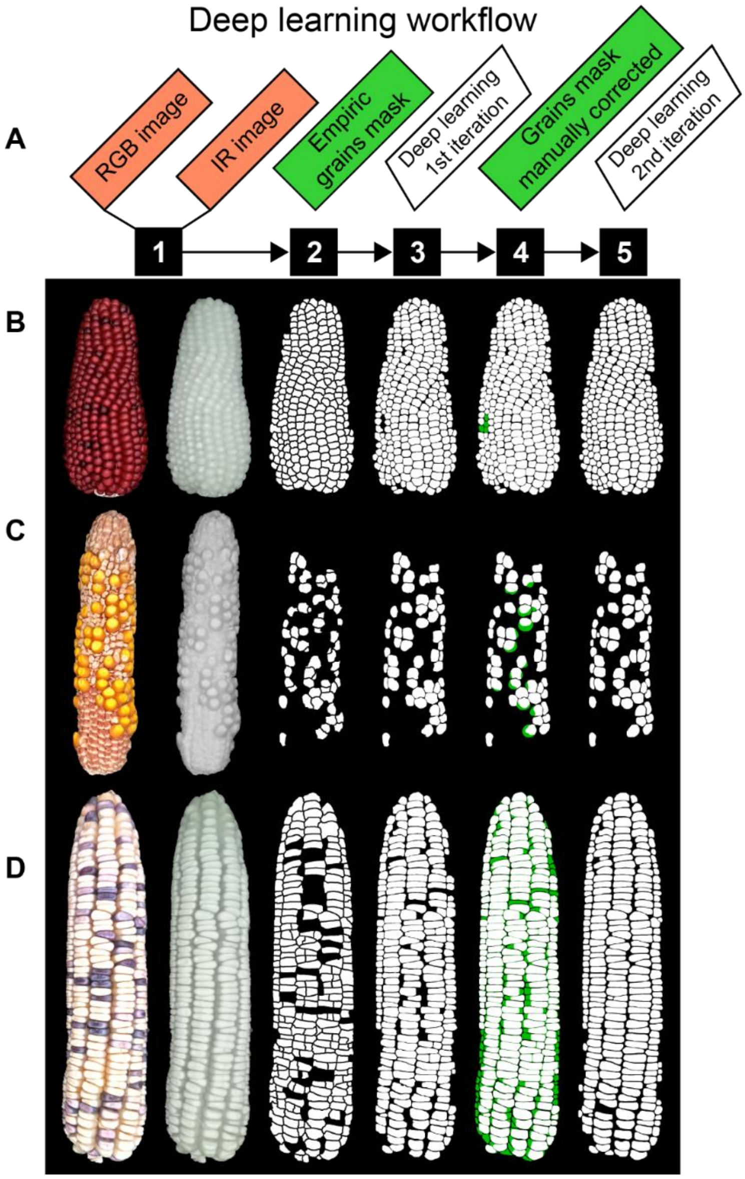


**Supplementary Fig. 2. Steps of the deep learning iteration workflow.** (A) Steps of the deep learning (DL) workflow and (B,C,D) example images used for 3 typical ears representing the problems encountered and the manual correction performed. (B) Typical ear with good empirical grain masks used directly for training the first iteration of deep learning (B2) and requiring minimum manual corrections (B3 to B4) to produce the routine segmentation method (B5). (C, D) Typical ears with false detections and erroneous grain segmentations from the empirical segmentation (C2, D2) and requiring major manual corrections (step 3 to 4) used for training the second deep learning iteration to produce the routine segmentation method (C5, D5). Orange boxes, image acquisition. White boxes, image, or data processing. Green boxes processed images or data. Green areas, manual grain mask corrections.

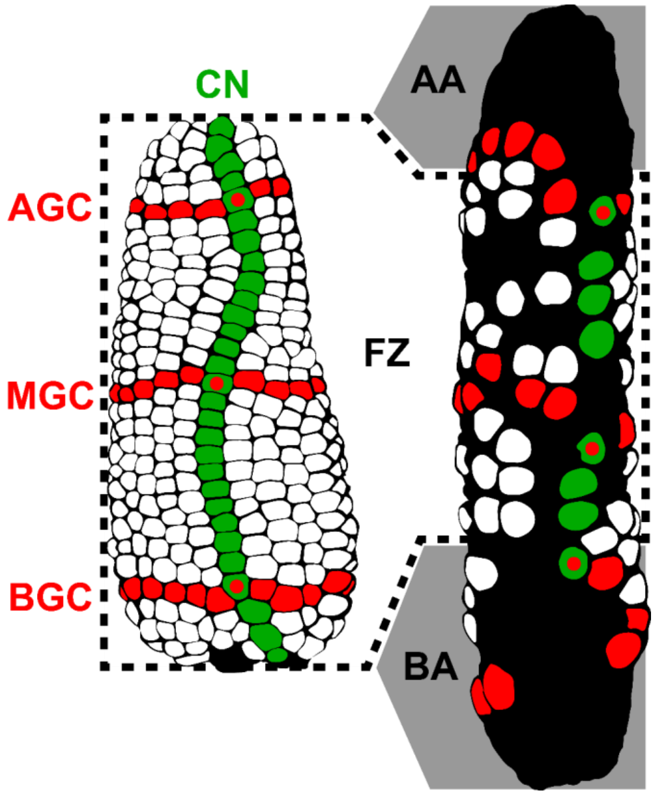


**Supplementary Fig. 3. Methodology for manual phenotyping of maize ears.** (A) Sample ear from the biological diversity panel. (B) Sample ear from the environmental diversity panel grown under water deficit conditions showing partial pollination and/or ovary and grain abortion. Red areas: example of grains used for manual phenotyping of basal (BGC), median (MGC) and apical (AGC) number of grains per cohort, particularly difficult to characterize on "scattered" ears. To standardize the measurement, the method consisted in counting the number of rows (= lines of grains along the ear perpendicular to the cohorts) with at least one grain per row for each third of the ear. Green areas: example of grains used for manual phenotyping of the average number of cohorts per ear (CN). The measurement was repeated 4 times around the ear. White zone: fertile zone of the ear (FZ), where the surface occupied by grains visually represents more than 50% of the visible surface of the ear. Grey zones: basal (BA) and apical (AA) abortion zones, where the surface occupied by grains visually represents less than 50% of the visible surface of the ear. The lengths of the FZ, BA, and AA zones were measured manually along the main axis of the ear.

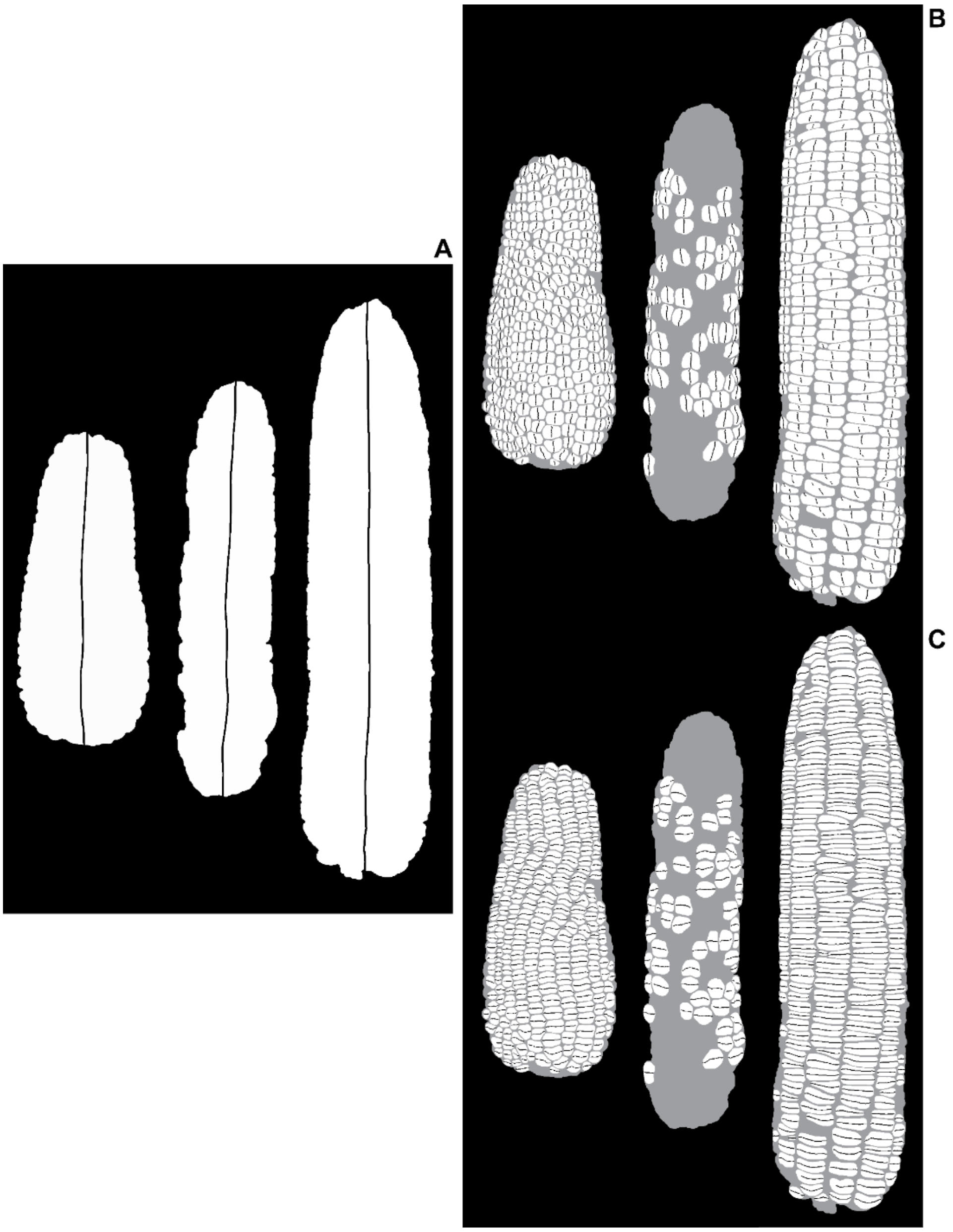


**Supplementary Fig. 4. (A) Illustration for 3 ears of the processing used to extract ear length from ear masks.** The black line represents the centerline of pixels along the main axis of the ear (vertical axis, starting from the bottom to the top of the ear), the measured ear length is the number of pixels in this line. (B) Illustration of the processing procedure to calculate the number of grains per cohort. Grain objects were reduced to a one-pixel wide vertical line along the principal axis. (C) Illustration of the processing procedure to calculate the number of grain cohorts. Grain objects were reduced to a one-pixel wide horizontal line along the ear axis perpendicular to the principal axis.

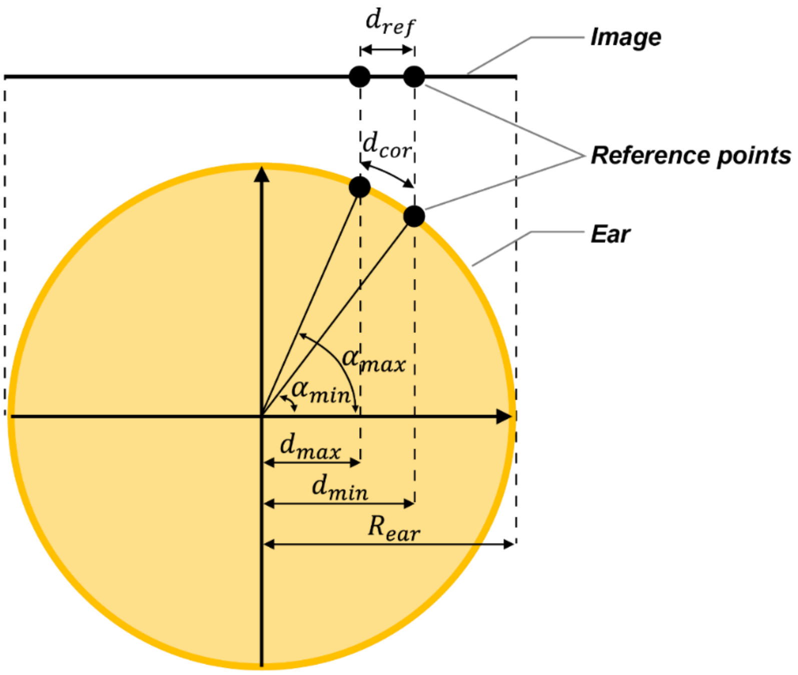


Corrected measurement: $d_{cor} = (\alpha_{max} - \alpha_{min}) * R_{ear}$

With: $R_{ear} = ear\ radius$, $\alpha_{max} = \cos^{-1}\left(\frac{d_{max}}{R_{ear}}\right)$ and $\alpha_{min} = \cos^{-1}\left(\frac{d_{min}}{R_{ear}}\right)$

**Supplementary Fig. 5. Illustration of the image correction applied to project the distances and positions of the reference points onto a hypothetical circular section of the ear.** Orange circle, boundaries of the ear. The reference image measurement ($d_{ref}$) is corrected using the horizontal distances measured between its extreme reference points (black dots) and the center of the ear ($d_{max}$ and $d_{min}$). The final corrected measured is an estimate of the length of the arc resulting from the projection of $d_{ref}$ onto a hypothetical perfect circle of radius ($R_{ear}$).

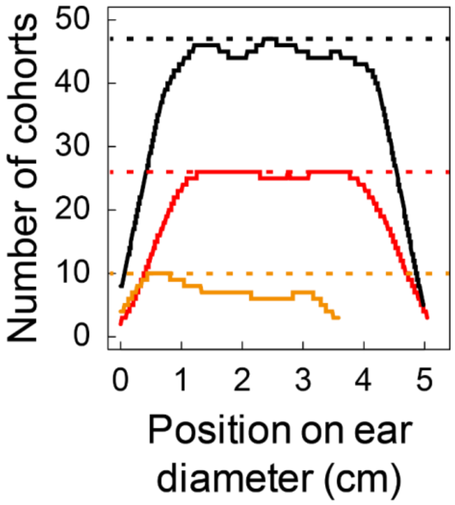


**Supplemental Fig. 6. Output of the estimation of the number of cohorts along the perpendicular ear axis.** For each position along the ear diameter (x), the number of cohorts (y) is calculated for each ear shown in Fig. 5 with the methodology presented in Supplementary Fig. S3. Black line, ear with white grains; red line, ear with vine grains; golden line, aborted ear with yellow grains. Dotted lines represent the maximum number of cohorts for each curve. The maximum number of cohorts is used as the output of the routine workflow and its average over the 6 ear sides is used for the correlation in Fig. 6. Each curve represents data from a single image/ear side.

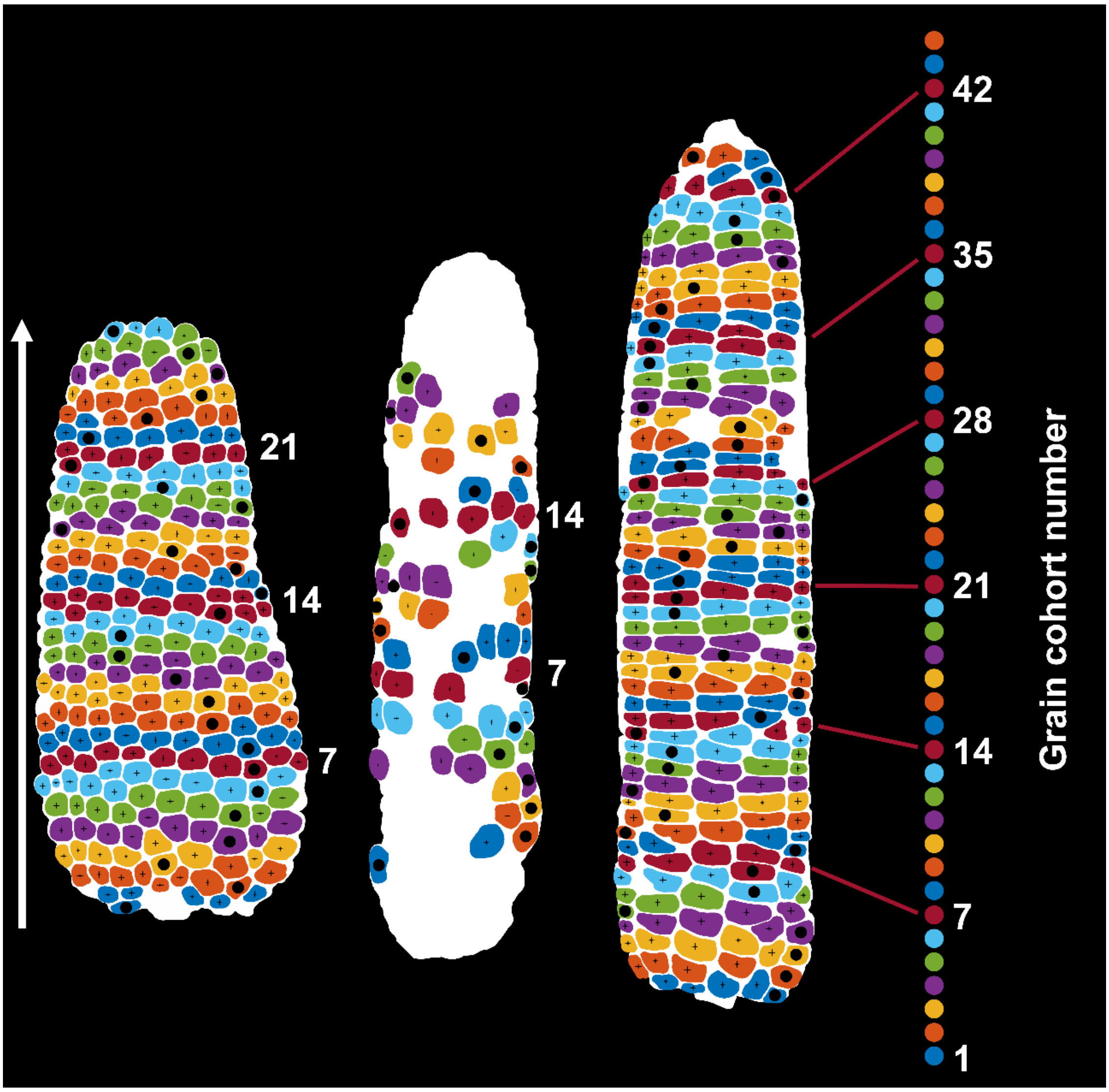


**Supplementary Fig. 7. Illustration of the cohort classification algorithm.** Grains are classified in cohorts (each identified by a color) by scanning the ear from bottom to top starting with the lowest grain (bold black dots on the barycenter). The classification is done sequentially, one cohort at a time. For each cohort, grains whose barycenter lie within a specific range of pixels along the main axis of the ear are classified into a common cohort, and removed for the rest of the classification process. Black dots: grain barycenter's. Colors: grains classified in the same cohort.

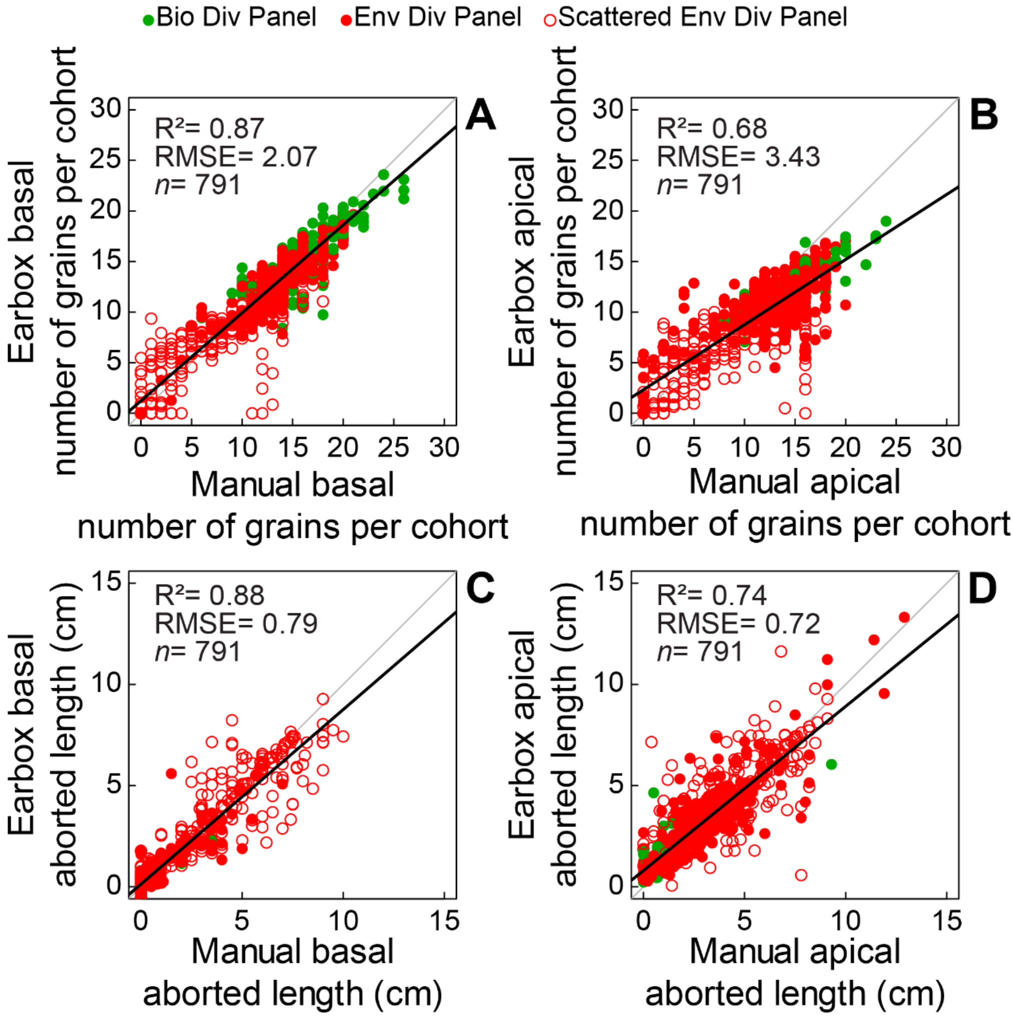


**Supplementary Fig. 8. Comparison of Earbox (y) and reference (x) data.** (A) Number of grains per cohort in the basal third. (B) Number of grains per cohort in the apical third. (C) Length of the basal aborted zone in centimeters. (D) Length of the apical aborted zone in centimeters. Green dots: ears from biological diversity panel; red dots: ears from environmental diversity panel. Empty red dots: scattered ears of the environmental diversity panel (Fig.3B). Grey line: bisector line. Black line: linear regression of data. $R^2$: correlation coefficient between x and y values, RMSE: root mean square error, *n*: number of observations in each graph.

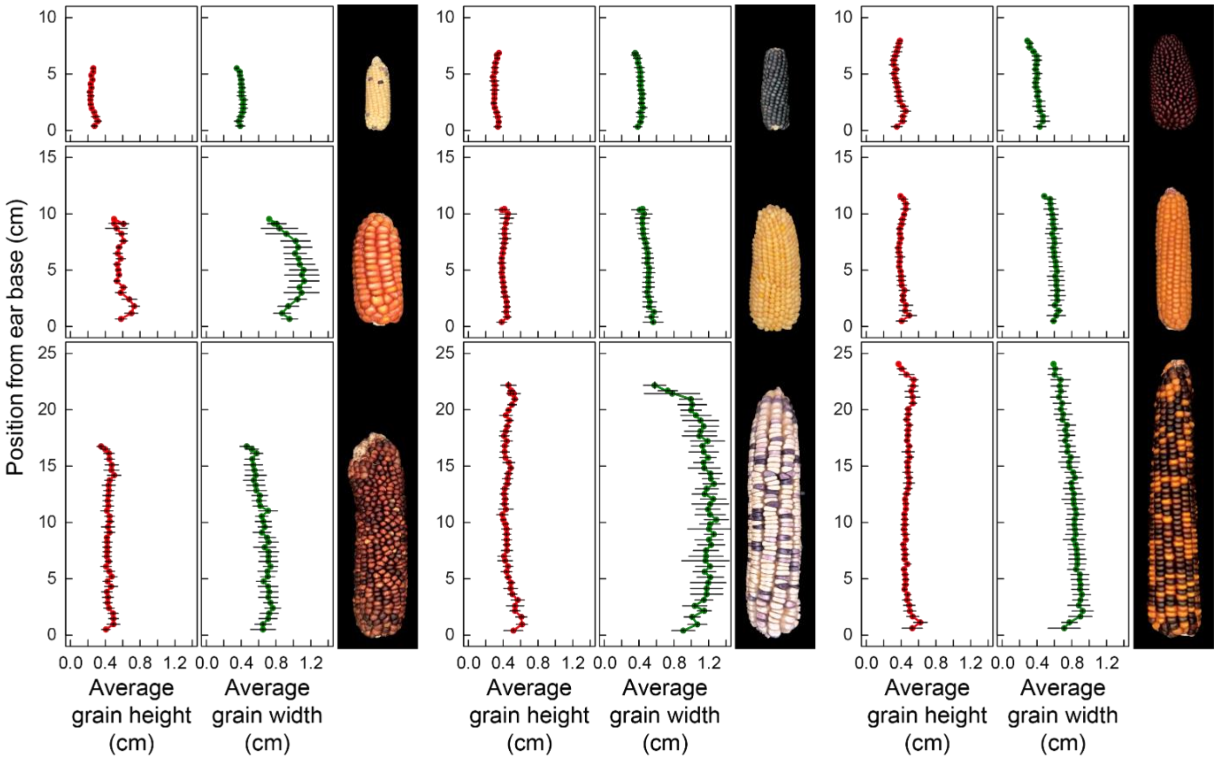


**Supplementary Fig. 9. Examples of grain dimensions as a function of cohort and position along the ear across 9 contrasting ears.** Each point represents the average dimension of all grains classified in the same cohort (Fig. 5P, 5Q, 5R) across the 6 sides (images) of the ear. Red dots, average grain height in centimeters. Green dots, average grain width in centimeters. Error bars, standard deviation. Errors for position from ear base are shown but are smaller than the dots.
